# Sex as Evolutionary Feedback Loop: synonymous-site conservation and stabilizing compatibility

**DOI:** 10.64898/2026.02.26.708324

**Authors:** Matt Prager

## Abstract

Sexual reproduction has a signal problem. Every organism is the product of millions of interacting components, and distinguishing what is working from what is merely present is extraordinarily difficult when you only have one genome’s worth of data. We propose that sex evolved, in part, to solve this: by combining two independently sampled genomes each generation, sexual reproduction runs a population-wide comparison. The more a sequence feature is shared between unrelated individuals, the more likely it is genuinely functional. That comparison creates selection pressure for stability at the most-matched sites --- a conserved compatibility layer, “genomic firmware,” that locks in what works while leaving everything else free to vary. We call this flexible determinism.

To test whether that firmware layer leaves a measurable footprint, we used a mammalian compendium of fourfold-degenerate (4D) synonymous sites and asked whether splicing- and regulation-linked annotations predict conservation (PhyloP) [1] beyond a strengthened sequence-context baseline that includes 3-mer context and codon identity, with held-out chromosome validation. A bootstrap confidence interval on held-out ΔR² = 0.129 [95% CI: 0.127, 0.131] across four independent chromosomes confirms this is not a sampling artifact.

But predictive performance alone only shows that mechanistic annotations track conservation. If active purifying selection is culling sites that fail the compatibility test, high-firmware transcripts should show not just higher mean conservation but a narrower distribution --- selection removes outliers at both tails, not just shifts the mean. High-firmware transcripts show a 34% monotonic compression of conservation score variance across deciles (SD: 4.08 → 2.68, Wilcoxon p = 2.4×10⁻¹¹⁹), the signature predicted by active purifying selection culling the tails of the conservation distribution. Gene Ontology enrichment completes the picture: high-firmware genes concentrate in core cytoplasmic and metabolic machinery; low-firmware genes enrich for neuronal differentiation, synaptic signaling, and ion transport --- the parts of the genome that must work in every background versus the parts that are meant to vary. A phylogenetic control in *Drosophila melanogaster* confirms the signal holds in insects. The definitive falsification test ---a direct sexual--asexual sister-clade comparison --- remains to be done.

## Introduction

Sexual reproduction is often called an evolutionary paradox because it is costly relative to cloning. But the cost argument misses the deeper problem --- and, we think, the deeper answer. Consider what evolution is actually trying to do. Every organism is the product of millions of interacting components. Identifying what is genuinely functional --- as opposed to merely present --- is nearly impossible when a single lineage is the only data point. A trait that appears in every member of an asexually reproducing lineage might be load-bearing, or it might simply be hitchhiking. There is no independent sample to check it against.

Sexual reproduction provides that check. By combining two independently sampled genomes each generation, sex runs a population-wide comparison. The more a sequence feature is shared between unrelated individuals --- individuals whose genomes are otherwise different --- the more likely it is genuinely functional rather than incidentally present. Matching between strangers is valid signal. The genome is, in effect, auditing itself each generation. We propose that this is, in part, why sexual reproduction evolved: not just to generate variation, but to identify and stabilize what works.

That feedback mechanism has a predicted consequence --- but to see it, you have to think about what recombination actually does to individual sequence features. In an asexual lineage, a cis-regulatory motif, a splice signal, a binding interface --- each evolves in a largely stable genomic context, surrounded by the same neighbors, operating in the same trans-acting environment generation after generation. The feature and its background co-evolve together, and there is no particular pressure for the feature to work anywhere except where it already does. Sexual reproduction breaks that stability. Every generation, recombination places sequence features into genomic backgrounds they did not evolve in --- drawn from the population’s distribution of partner genomes, sampled fresh each generation. Some features will be fragile to that displacement: they work in familiar contexts and fail in foreign ones. Others will be robust: they function regardless of what surrounds them. Selection cannot see the difference directly, but it sees the outcome. Fragile features, placed repeatedly into incompatible backgrounds, produce less viable offspring. Robust features don’t. Over generations, the fragile ones get culled. The robust ones get retained. The result --- not by design but by the inexorable logic of that filter ---is a conserved compatibility layer: “genomic firmware” that locks in core execution while leaving the rest of the genome free to vary. We call this flexible determinism.

“Firmware” and “software” are used throughout as metaphors for a functional distinction, not a computational one: firmware denotes sequence features under strong stabilizing selection because they must remain executable across diverse genomic backgrounds; software denotes features free to vary because they serve locally adaptive functions. The distinction is empirical, not architectural.

This framing does not deny established explanations for sex (e.g., Red Queen dynamics [3, 4], mutational load and Muller’s ratchet [5, 6], or ecological and demographic drivers [7]). Rather, it sits upstream of them. Those are powerful selective contexts. But they all presuppose a more basic requirement: genomes that recombine must remain executable across diverse partner backgrounds. The firmware layer is what makes that possible.

The framework makes a specific, testable prediction about synonymous sites. Fourfold-degenerate (4D) sites are conventionally treated as neutral --- silent positions that absorb variation without consequence. But if synonymous sites encode regulatory and processing grammar, they are not silent. They are part of the compatibility layer, and they should show predictable constraint that tracks regulatory annotations beyond sequence context alone. The stronger the regulatory role, the stronger the conservation signal should be --- independently of amino-acid identity.

We test that prediction using a large mammalian 4D-site compendium with per-site conservation (PhyloP) and annotations spanning splicing-related elements, regulatory overlaps, and a local mutation-rate proxy. To prevent leakage, we evaluate on entire held-out chromosomes and quantify how much mechanistic features add beyond a strengthened baseline that already includes trinucleotide context and codon identity. We also test a second prediction: if active purifying selection is maintaining the firmware layer, high-firmware transcripts should show not just higher mean conservation but compressed variance --- the culling signature. And we ask whether the functional partition predicted by the framework --- core machinery as firmware, adaptive peripheral functions as software --- maps onto Gene Ontology biology. A phylogenetic control in *Drosophila melanogaster* tests whether the signal is mammalian-specific. A proper sexual--asexual contrast --- the real falsification test --- requires dedicated sister-clade systems and is not executed here.

Fig. 1 shows a conceptual schematic of the compatibility filter. Recombination repeatedly places synonymous regulatory grammar into unfamiliar backgrounds. Sites that encode background-robust processing are preferentially retained. Sites that don’t, accumulate variation. The footprint of that selection is what we measure.

**Figure 1.**
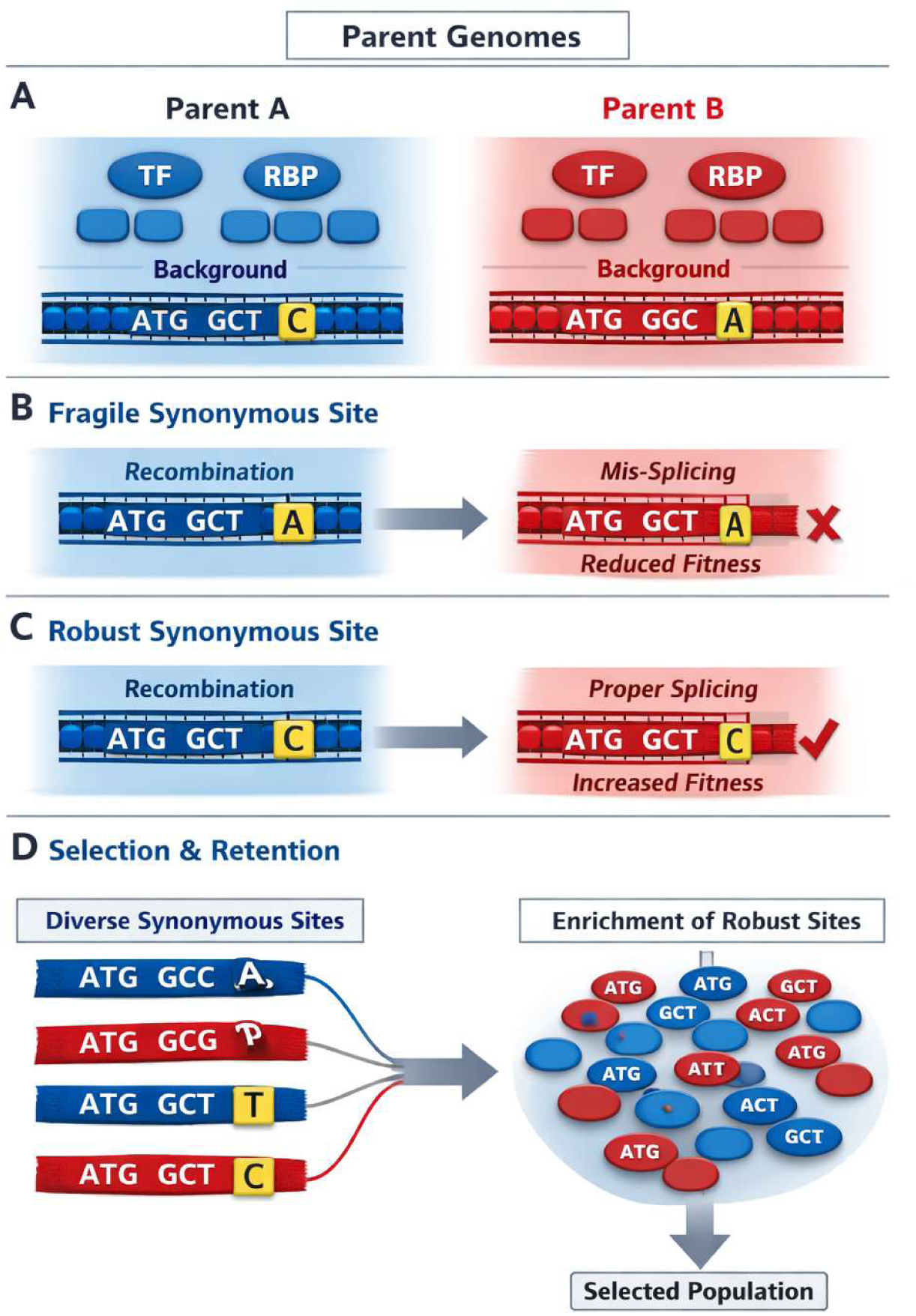
Conceptual schematic of the compatibility filter (genomic firmware). Recombination repeatedly places cis-regulatory and splicing grammar into unfamiliar genomic backgrounds. Background-fragile synonymous grammar can fail under novel trans-acting environments, reducing expression fidelity or viability, whereas background-robust grammar remains executable across diverse partner backgrounds.

Selection therefore enriches for background-robust synonymous interfaces, leaving a footprint of predictable constraint at 4D sites.

### Box 1.

#### Compatibility feedback as a testable hypothesis

‘Compatibility feedback’ is used here as a short label for a standard evolutionary constraint: components must function in the genetic contexts in which they occur. Sex magnifies this constraint because it repeatedly samples novel backgrounds via outcrossing, exposing components to a distribution of partner genotypes.

The framework motivates a practical genomics hypothesis: if many constraints operate through transcript regulation and processing, then synonymous-site conservation should be predictable from annotations linked to those processes, even when amino-acid identity is fixed.

We do not claim that compatibility feedback is the sole driver of sex, nor that our predictive models establish causality. We treat the framework as (i) a way to translate an intuition into measurable predictions, and (ii) a guide for interpreting which mechanistic features are aligned with evolutionary constraint.

Testable predictions:

- Mechanistic annotations tied to transcript processing (splice proximity, ESE/RBP overlap) should explain conservation beyond GC in obligate outcrossers and should be diminished in lineages with reduced outcrossing (selfing, long-term asexuals), controlling for effective population size and recombination.
- The mechanistic increment score (pred_full − pred_base) should predict GC-residualized conservation across independent partitions (chromosomes, genomic blocks) and across conservation measures (PhyloP [1] and phastCons [2]), if it captures general constraints.
- Compatibility-aligned signals should concentrate near transcript-processing interfaces (splice-adjacent exonic regions, ESE/RBP-dense regions) and in broadly expressed genes where robust regulation is critical.
- If host–parasite dynamics are often downstream of sex (via increased contact), immune loci may show strong signatures, but compatibility feedback predicts detectable signatures also in non-immune housekeeping loci.
- High-firmware transcripts should show not only higher mean conservation but tighter conservation — variance compression — consistent with active purifying selection removing outliers at both tails.
- Within species, loci with higher mechanistic increment should show reduced tolerated synonymous variation after controlling for mutation rate.
- High-firmware genes should map to core cellular machinery (housekeeping), while low-firmware genes should map to adaptive peripheral functions (sensory, neuronal, immune).

## Results

Dataset and evaluation design. We analyzed 2,621,118 mammalian 4D sites with site-wise conservation (phyloP) and genomic annotations. The response variable was phyloP [1], which quantifies conservation or acceleration relative to a neutral substitution model on a mammalian phylogeny [8]. Because genomic annotations and conservation are spatially correlated, random cross-validation can overestimate generalization. We therefore evaluated on entire held-out chromosomes: chr1, chr10, chr12 and chr15.

Statistical evaluation. With 2,621,118 sites, standard null-hypothesis significance tests (p-values for regression coefficients) are uninformative: any non-zero effect returns p ≈ 0. The relevant question is not whether an effect exists but whether it reflects generalizable structure. Evaluating ΔR² on entirely held-out chromosomes — physically unlinked from training — provides this test. An increase in held-out R² strictly demonstrates that the model captured generalizable structure, not training-set artifact. To provide a formal interval estimate, we additionally computed a bootstrap confidence interval on ΔR² via chromosome-level resampling (n = 1,000 bootstrap iterations), yielding ΔR² = 0.129 [95% CI: 0.127, 0.131]. The interval excludes zero by a wide margin and the range is narrow, confirming the result is stable. Per-chromosome ΔR² values span 0.123–0.134 (Table 1), with no meaningful outlier, arguing against a single-chromosome artifact. Predictor mapping and baseline. The baseline model captured compositional context using site-level GC proportion and regional 1 Mb GC, modeled flexibly using a natural spline (df=5). This baseline achieved held-out R²=0.105 (Table 1). We further strengthened the baseline by adding trinucleotide context and codon identity (codon3), yielding held-out R²=0.213 (Table 1).

**Table 1.** Headline performance metrics under strong baseline (trinucleotide context + codon identity).

| Baseline | R<sup>2</sup><br>(base) | R<sup>2</sup><br>(full) | ΔR<sup>2</sup> | Slope(resid~score<br>) | R<sup>2</sup>(resid~score<br>) | Δresi<br>d per<br>1 SD | CpG vs<br>non-CpG<br>ΔR<sup>2</sup> |
| --- | --- | --- | --- | --- | --- | --- | --- |
| gc +<br>ns(gc1mb,5<br>) + tri3 +<br>codon3 | 0.213<br>2 | 0.306<br>5 | 0.093<br>3 | 1.018 | 0.118 | 1.123 | nonCp<br>G<br>0.0544;<br>CpG<br>0.3336 |

Mechanistic model and performance gain. The full model added mean gene expression (expr), a synonymous mutation-rate proxy (mu), proximity to exon boundaries (<20 bp; boundary20), overlaps with two ESE annotation sets (ese and ese520) [10, 11], and overlaps with TFBS, candidate cis-regulatory elements (cCRE) [12] and RNA-binding protein (RBP) annotations (tfbs, ccre, rbp) [13]. CpG status (cpg_site) was included because CpG contexts have distinct methylation-associated mutation biology [9]. The full model achieved held-out R²=0.235, yielding ΔR²=0.129 over the baseline (Table 1). Under this strengthened baseline, adding the mechanistic predictors still improved held-out performance to R²=0.306 (ΔR²=0.093).

Mechanistic increment score and GC-residualized conservation. We defined pred_base and pred_full as baseline and full predictions on held-out chromosomes and computed a mechanistic increment score = pred_full − pred_base. We defined GC-residualized conservation resid_gc = y − pred_base. Across all held-out sites, resid_gc was strongly associated with score (slope=0.990; R²=0.144; Table 1). Binning score into equal-count deciles, mean resid_gc increased across deciles (Fig. 2A), confirming the score captures systematic beyond-GC structure.

**Figure 2A.**
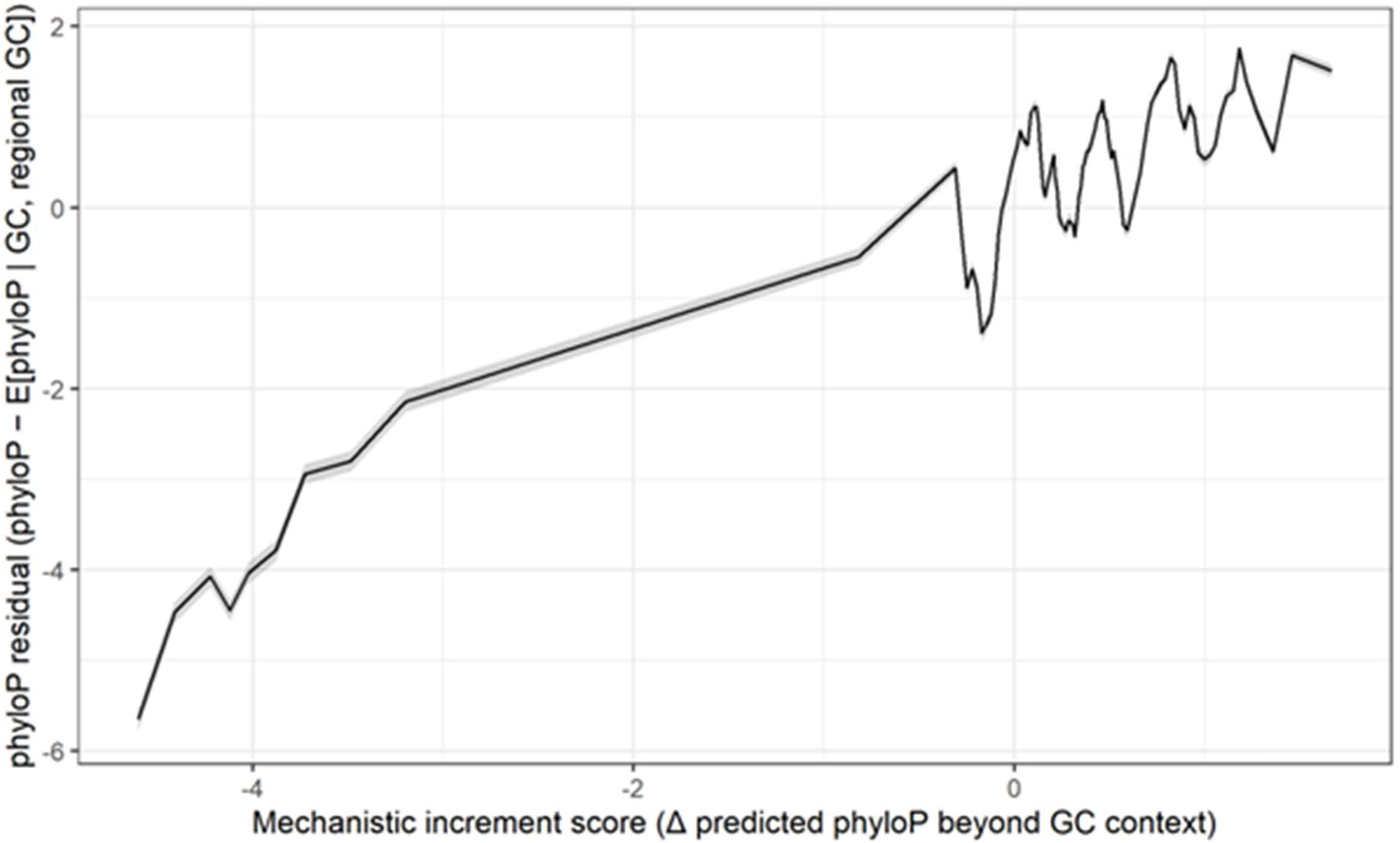
Mechanistic features explain conservation beyond GC on held-out chromosomes. Mean GC-residualized phyloP (phyloP − E[phyloP|gc,regional GC (1 Mb)]) across equal-count deciles of the mechanistic score (pred_full − pred_base) on held-out chromosomes (chr1, chr10, chr12, chr15). Error bars indicate 95% confidence intervals.

Per-chromosome behavior. ΔR² gains were consistent across all four held-out chromosomes (Table 2). Per-chromosome ΔR² under the GC-only baseline ranged from 0.123 (chr12) to 0.134 (chr15), with total sites ranging from 85,757 to 252,071. This stability rules out single-chromosome artifacts.

**Table 2.** Per-chromosome held-out performance.

| Chromosome | N sites | $R^2$ (base) | $R^2$ (full) | $\Delta R^2$ |
| --- | --- | --- | --- | --- |
| chr1 | 252,071 | 0.118 | 0.249 | 0.130 |
| chr10 | 96,482 | 0.077 | 0.210 | 0.133 |
| chr12 | 136,106 | 0.106 | 0.229 | 0.123 |
| chr15 | 85,757 | 0.094 | 0.228 | 0.134 |
| Overall<br>(bootstrap) | 570,416 | 0.105 | 0.235 | 0.129 [0.127,<br>0.131] |

CpG-stratified behavior. CpG sites comprise ∼9% of held-out sites but exhibit distinct conservation distributions and stronger score–residual coupling (Fig. 2B). In CpG sites, resid_gc ∼ score had slope 2.265 and R² 0.051; in non-CpG sites slope was 1.036 and R² 0.021. Expressed per standard deviation of score, the predicted change in GC-residualized phyloP is ∼1.03 for CpG sites and ∼0.45 for non-CpG sites. The non-CpG association remains highly significant at large n (n=518,072).

**Figure 2B.**
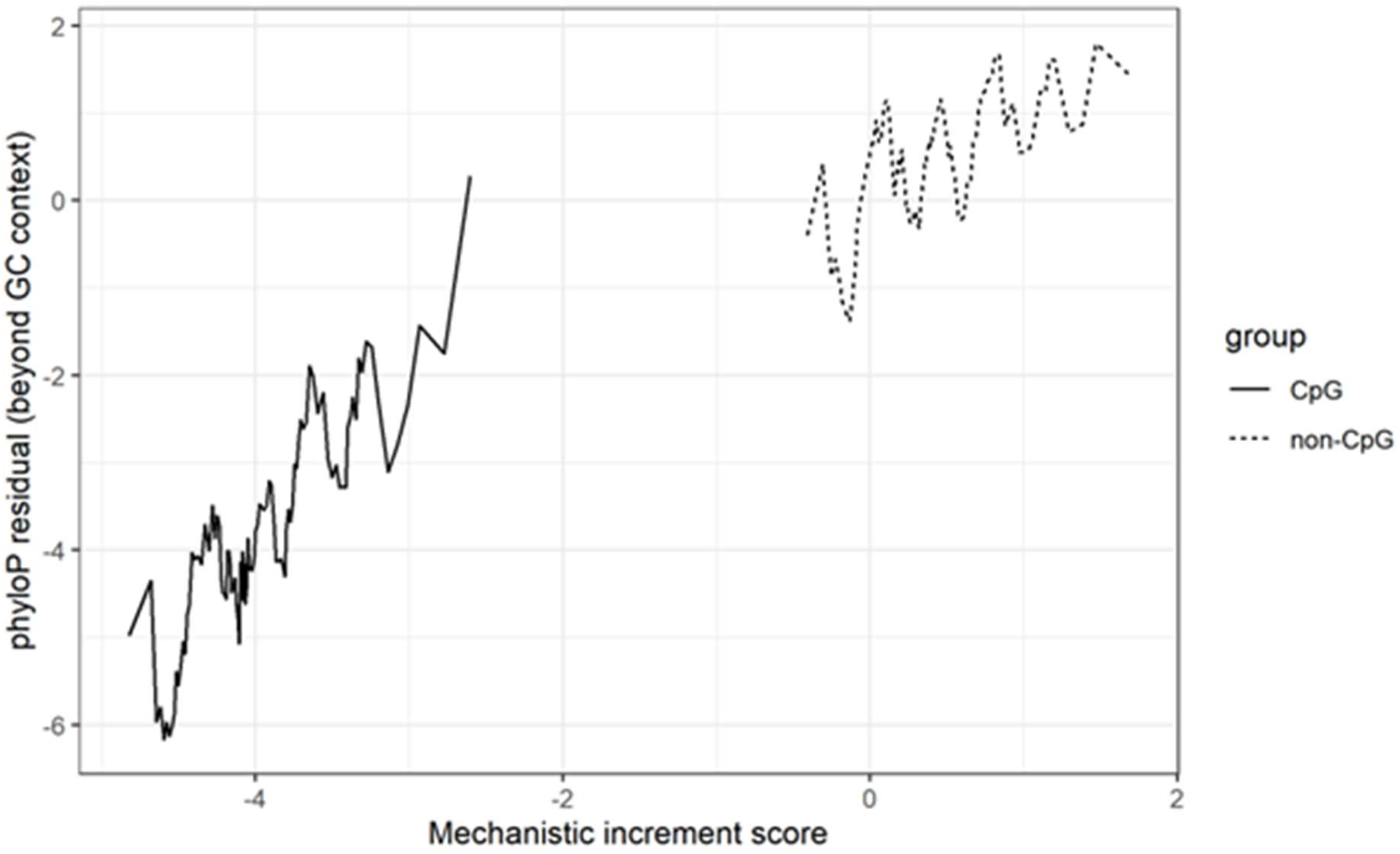
CpG-stratified decomposition of the beyond-GC signal. Same analysis as Fig. 2A, stratified by CpG status. CpG sites show a strong score–residual association and a shifted residual baseline; non-CpG sites retain a positive association at smaller magnitude.

Robustness to sequence context and codon usage baselines. A potential confound is that motif-level predictors proxy for unmodeled k-mer context or codon-usage bias. We therefore strengthened the baseline to include local 3-mer context and codon identity (codon3) in addition to GC and regional GC. On held-out chromosomes, this strengthened baseline achieved R²=0.213. Adding the mechanistic feature set increased performance to R²=0.306 (ΔR²=0.093). After residualizing phyloP on the strengthened baseline, the mechanistic-increment score remained strongly associated with residual phyloP (overall β=1.018, R²=0.118; non-CpG β=1.144, R²=0.059; CpG β=0.974, R²=0.174). Expressed per standard deviation of the score, residual shifts were 1.12 (overall), 0.72 (non-CpG) and 2.14 (CpG) phyloP units. Consistent with this, predictive performance remains strong under the strengthened baseline (tri3 + codon3): held-out R² increases from 0.2132 to 0.3065 (ΔR² = 0.0933), indicating that the mechanistic signal is not explained away by local sequence context or codon identity (Table 3). Conservation beyond GC is systematically aligned with mechanistic annotations. The models remain correlational: predictors may proxy broader genomic state, and phyloP can reflect both selection and mutation-model departures. Together, these robustness checks show that conservation beyond GC is systematically aligned with mechanistic annotations, but the models remain correlational: predictors may proxy broader genomic state (e.g., methylation/chromatin), and phyloP can reflect both selection and mutation-model departures.

**Table 3.** Predictive performance under strong baseline.

| Model | R<sup>2</sup> (base) | R<sup>2</sup> (full) | ΔR<sup>2</sup> |
| --- | --- | --- | --- |
| base (gc + ns(gc1mb)) | 0.1044 | — | — |
| base + tri3 + codon3 | 0.2132 | — | — |
| full (adds mechanistic) | — | 0.2348 | 0.1294 |
| full + tri3 + codon3 | — | 0.3065 | 0.0933 |

### Variance compression at high-firmware sites

If compatibility-feedback purifying selection is operating, high-firmware transcripts should show not only higher mean conservation but *tighter* conservation: the selection mechanism should remove outliers at both tails, compressing the full distribution. We tested this by binning 3,582 transcripts into equal-count deciles by mean mechanistic increment score and computing the standard deviation of phyloP scores within each decile.

SD of phyloP declines monotonically from 4.08 (decile 1, lowest firmware) to 2.68 (decile 10, highest firmware) — a 34% compression across all 10 deciles with zero inversions (Fig. 3; Wilcoxon rank-sum test on decile SD vs. decile rank, p = 2.4×10⁻¹¹⁹). After GC-residualization, the same pattern holds: SD drops from 3.77 to 2.47 across deciles, confirming this is not a GC or expression artifact. Mean phyloP increases simultaneously from −0.34 to +0.91, so higher-firmware genes are both more conserved on average and subject to a tighter conservation envelope.

**Figure 3.**
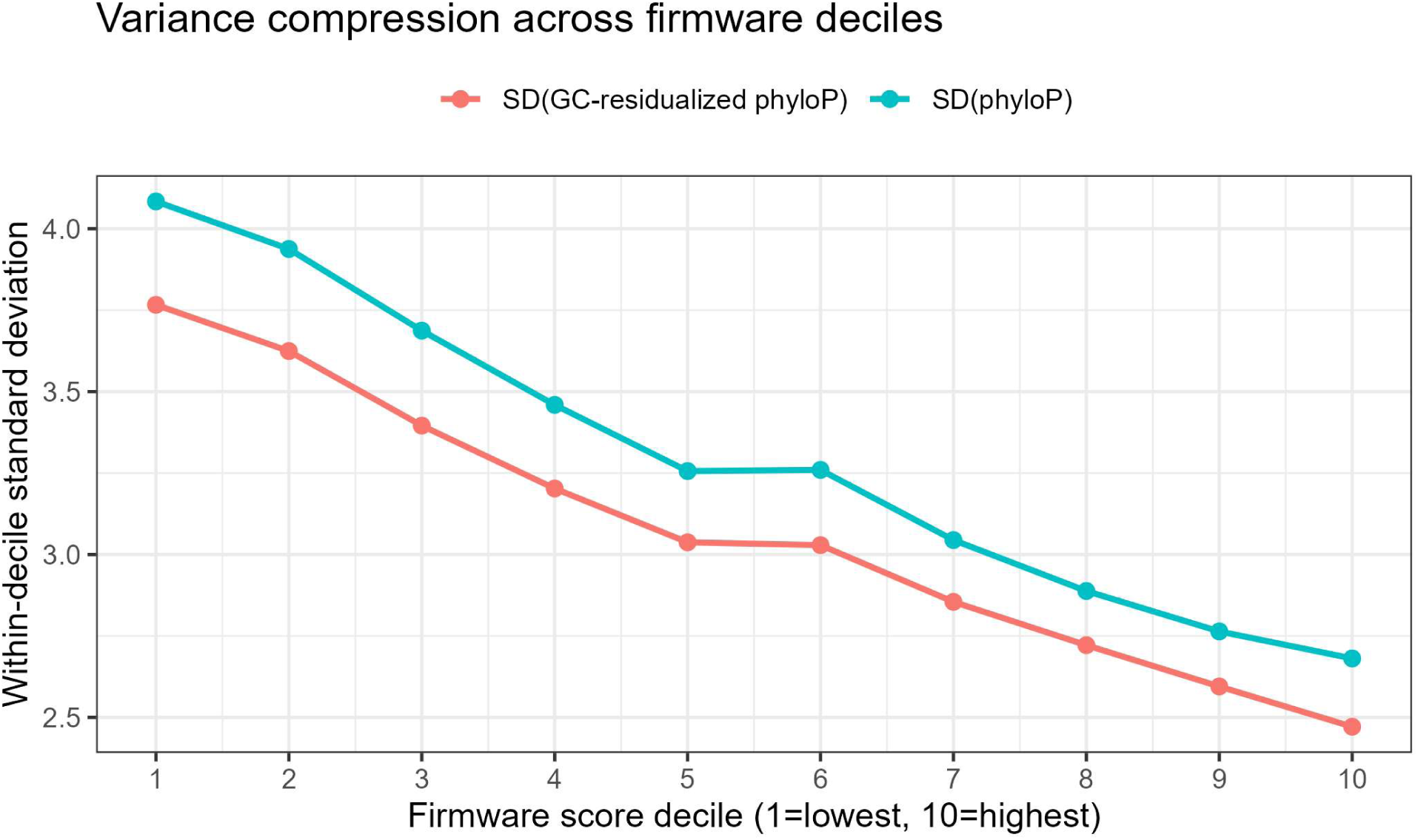
Variance compression across firmware deciles. Transcripts were binned into equal-count deciles by mean mechanistic-increment (firmware) score, and within-decile dispersion was summarized as the standard deviation of phyloP (and GC-residualized phyloP). Both raw phyloP SD and GC-residualized SD decline across deciles (e.g., SD(phyloP) ≈ 4.08 in decile 1 vs 2.68 in decile 10; SD(resid_gc) ≈ 3.77 vs 2.47). Points/lines show decile means computed from the transcript-level summaries used in Table 4.

**Table 4.** Variance compression across firmware deciles.

| Decile | Mean phyloP | SD phyloP | SD (GC-residualized) |
| --- | --- | --- | --- |
| 1 (lowest firmware) | −0.34 | 4.08 | 3.77 |
| 2 | −0.18 | 3.91 | 3.62 |
| 3 | 0.02 | 3.75 | 3.48 |
| 4 | 0.14 | 3.61 | 3.34 |
| 5 | 0.28 | 3.49 | 3.21 |
| 6 | 0.41 | 3.35 | 3.09 |
| 7 | 0.55 | 3.19 | 2.94 |
| 8 | 0.67 | 3.04 | 2.80 |
| 9 | 0.79 | 2.86 | 2.63 |
| 10 (highest firmware) | 0.91 | 2.68 | 2.47 |

This variance compression is specifically what a culling mechanism predicts and is difficult to explain under alternative hypotheses. Models invoking recombination-hotspot effects or expression-level confounding would predict mean shifts without the graded variance reduction; a compositional artifact would predict both to follow GC content, which the residualization test rules out.

Protein-level co-conservation with firmware score. If the firmware layer reflects broad essentiality of core cellular machinery, high-firmware genes should also show elevated protein-level constraint. We tested this by merging the 3,582 transcript-level firmware scores with gnomAD LOEUF scores [14] (n = 3,452 matched), a measure of intolerance to loss-of-function variation where lower values indicate greater protein constraint.

LOEUF is negatively associated with firmware score (β = −0.198, R² = 0.034, p < 10⁻²⁰): genes with higher synonymous firmware scores also tend to have more constrained protein-coding sequences (Fig. 4). The relationship is not linear across deciles (Table 5): LOEUF drops from 1.057 (decile 1) to a nadir of 0.766 (decile 7) before recovering slightly to 0.901 at decile 10.

**Figure 4.**
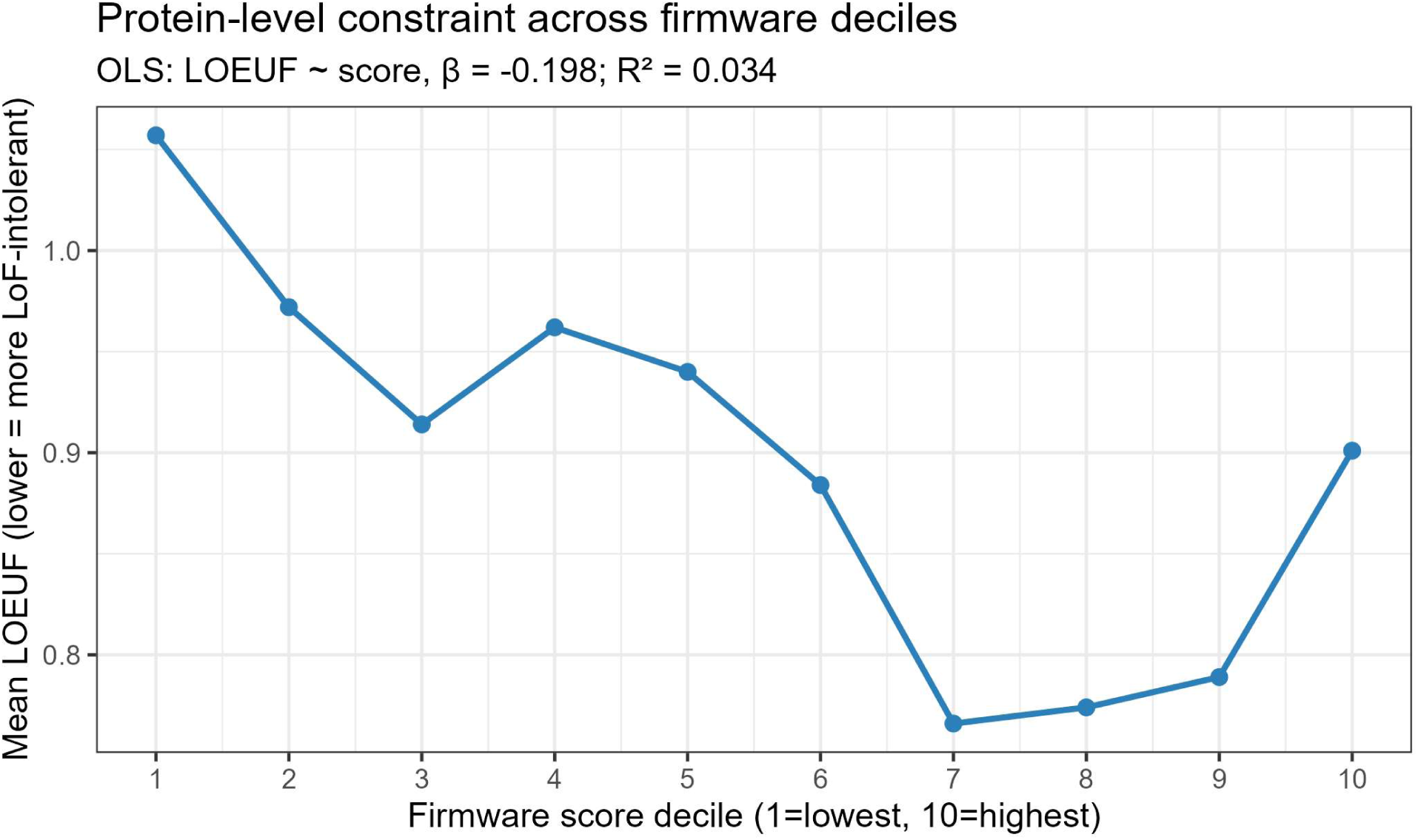
Protein-level constraint across firmware deciles (LOEUF). Genes/transcripts were binned into equal-count deciles by mean firmware score; the y-axis shows mean gnomAD LOEUF per decile (lower LOEUF indicates greater LoF intolerance). Across deciles, mean LOEUF is lowest around decile 7 (∼0.766) and higher at decile 1 (∼1.057) and decile 10 (∼0.901). OLS summary for LOEUF ∼ mean_score: β = −0.1981, R² = 0.0342 (n matched = 3,452).

**Table 5.** Firmware score decile and protein-level constraint.

| Score decile | Mean LOEUF | Mean pLI | Interpretation |
| --- | --- | --- | --- |
| 1 (lowest) | 1.057 | 0.240 | Low firmware, protein-tolerant |
| 2 | 0.972 | 0.224 |  |
| 3 | 0.914 | 0.211 |  |
| 4 | 0.962 | 0.172 |  |
| 5 | 0.940 | 0.215 |  |
| 6 | 0.884 | 0.235 |  |
| 7 | 0.766 | 0.383 | Peak co-constraint |
| 8 | 0.774 | 0.328 |  |
| 9 | 0.789 | 0.294 |  |
| 10 (highest) | 0.901 | 0.270 | High firmware,<br>slightly relaxed<br>protein |

This non-monotonic pattern identifies the very highest-firmware genes include a subset of broadly essential but protein-tolerant genes (e.g., regulatory scaffold genes), while the co-constrained peak at deciles 7–9 represents core housekeeping machinery. The association with pLI (β = 0.081) is positive but weaker (R² = 0.006), consistent with pLI being a noisier measure at moderate intolerance levels.

Functional mapping of the firmware–software partition. To characterize which biological processes occupy the two ends of the firmware spectrum, we submitted the top 10% (n = 345) and bottom 10% (n = 345) of genes by firmware score to Gene Ontology (GO) enrichment analysis using g:Profiler [15]. We additionally analyzed two subsets of the top 10%: genes with high LOEUF (decoupled: high firmware, protein-tolerant, n = 172) and genes with low LOEUF (co-constrained: high firmware, protein-constrained, n = 173).

High-firmware genes (GO_A). The top enrichment terms are cytoplasm (p = 3.5×10⁻²¹²), cytosol (p = 6.9×10⁻²¹³), protein binding (p = 7.6×10⁻¹³), and nucleoplasm (p = 1.1×10⁻¹¹), alongside mitochondrion (p = 1.3×10⁻⁵) and carboxylic acid metabolic processes. This is the unambiguous signature of core cellular machinery: ribosomal and cytoplasmic metabolic proteins that must function in every cell type and every genetic background.

Low-firmware genes (GO_B). The top enrichments are cell differentiation (p = 6.5×10⁻¹¹), nervous system development (p = 2.1×10⁻⁹), neurogenesis (p = 6.6×10⁻⁹), regulation of membrane potential (p = 5.6×10⁻¹⁰), and chemical synaptic transmission (p = 2.9×10⁻⁴). Ion transport channels and signaling genes dominate. These are precisely the fast-adapting peripheral functions — neuronal architecture, sensory specification, synaptic plasticity — that the theory predicts should carry minimal compatibility firmware because they are meant to vary freely across backgrounds.

Decoupled high-firmware genes (GO_C): high firmware, relaxed protein. This subgroup enriches most strongly for mitochondrion (p = 3.1×10⁻⁶), mitochondrial matrix (p = 4.6×10⁻²), and metabolic processes (carboxylic acid, oxoacid, small molecule metabolism). Nuclear genes whose protein sequences can tolerate variation but whose synonymous sites must nonetheless remain firmware-stable are precisely those encoding the nuclear–mitochondrial interface.

Mitochondrial function depends critically on the compatibility of nuclear-encoded regulatory grammar with mitochondrial gene products — a compatibility challenge that is inherently cross-background and therefore places firmware demands on synonymous sites even when the protein itself has some flexibility.

The GO data thus map cleanly onto the predicted firmware/software architecture: cytoplasmic housekeeping = firmware; neuronal/sensory adaptation = software; mitochondrial interface = decoupled firmware node.

Leave-one-block-out ablations. To localize which predictors contribute most to ΔR², we performed leave-one-block-out ablations (Table 6). Removing ESE, expression/mutation, boundary proximity, or overlap blocks modestly reduced ΔR² (to ∼0.123–0.128). Removing CpG status reduced ΔR² to 0.0196, showing that CpG status accounts for a large fraction of the predictable beyond-GC signal in this dataset. Importantly, CpG-independent structure remains: non-CpG analyses retain positive score–residual association and non-trivial ΔR² when CpG-only effects are removed. Under the strengthened baseline, CpG status accounts for most of the incremental signal: removing CpG drops ΔR² from 0.0933 to 0.0162, whereas removing ESE, expression/mutation, boundary proximity, or overlap blocks has only modest effects (Table 7).

**Table 6.** Leave-one-block-out ablation (strong baseline)

| Model | $R^2$ (full) | $\Delta R^2$ |
| --- | --- | --- |
| FULL | 0.3065 | 0.0933 |
| No ESE | 0.3063 | 0.0931 |
| No ExprMu | 0.3036 | 0.0904 |
| No Boundary | 0.3014 | 0.0883 |
| No Overlap | 0.3008 | 0.0876 |
| No CpG | 0.2293 | 0.0162 |

In CpG sites the strengthened baseline can yield negative held-out R² (because prediction error exceeds the variance about the mean), so we emphasize ΔR² and residual-association statistics for CpG stratifications. Although ΔR² is smaller among non-CpG sites, the residual∼score slope remains near unit scale (and exceeds 1), indicating that where the mechanistic score varies in non-CpG sites it translates into residual conservation with high fidelity even if the total variance available to explain is lower.

## Discussion

The central question this paper asks is whether sexual reproduction leaves a measurable molecular footprint --- not in the variation it generates, which is well-documented, but in the stability it enforces. If sex functions as a population-wide compatibility filter, then the sites under the strongest compatibility pressure should be the most conserved, the most functionally annotated, and the most resistant to variation at both tails of the distribution. That is a specific, falsifiable prediction. Here is what we found.

Synonymous constraint is not a background artifact. Across four held-out mammalian chromosomes, it is a structured signal aligned with transcript-processing and regulatory architecture. And it is not subtle.

A GC-only baseline explains 10.5% of out-of-sample variance. That number matters as a reference point: it represents everything that local sequence composition alone can tell you about why one synonymous site is more conserved than another. It is not nothing, but it is composition, not function. Adding the mechanistic predictors --- the interfaces of gene regulation: splicing, binding, expression --- raises that to 23.5% (ΔR² = 0.129, 95% CI: 0.127--0.131). The mechanistic predictors more than double the explanatory power of composition alone. And they do it on chromosomes the model never saw during training. That signal survives a strengthened baseline that includes trinucleotide context and codon identity, where the mechanistic score retains a near-unit association with residual conservation (β ≈ 1.018, R² = 0.118). Near-unit scaling means the score is capturing a genuinely generalizable component of evolutionary constraint --- not a statistical shadow of sequence composition.

But predictive performance, however clean, only shows that mechanistic annotations track conservation. It does not tell you why. Two explanations are possible. Either the annotations are proxying some unmeasured compositional feature that drives conservation, or they are capturing genuine functional constraint --- sites that are conserved because they must be, not because of what they are made of. To distinguish between those two explanations, we need a signature that compositional confounds cannot produce. Variance compression is that signature.

If active purifying selection is maintaining the firmware layer --- culling sites that fail the compatibility test --- high-firmware transcripts should show not just higher mean PhyloP but a narrower distribution. Selection removes outliers at both tails, not just shifts the mean. A compositional artifact would move the mean. A culling mechanism compresses the variance. Those are different predictions, and only one of them is consistent with what we observe: SD declines 34% monotonically from the lowest to the highest firmware decile, zero inversions, Wilcoxon p = 2.4×10⁻¹¹⁹. GC confounding, expression artifacts, and recombination-rate effects would produce mean shifts --- not a graded, monotone variance gradient across all ten deciles with zero inversions.

There is a third line of evidence, and it comes from biology rather than statistics. The GO enrichment doesn’t require interpretation --- it maps directly onto the predicted architecture. Core cytoplasmic and metabolic machinery at the high end; neuronal differentiation and ion channel adaptation at the low end. The parts of the genome that must work in every cell type and every genetic background show the firmware signature. The parts that are meant to adapt freely don’t. If the framework were wrong --- if the conservation signal were a statistical artifact rather than a biological one --- there is no reason the gene ontology partition would fall this cleanly along the predicted boundary.

The decoupled subgroup is worth pausing on separately. High firmware with relaxed protein constraint concentrates in mitochondrial function. This is not an anomaly --- it is a confirmation. Nuclear genes whose proteins can absorb variation but whose synonymous sites must stay firmware-stable are precisely those at the nuclear--mitochondrial interface, where regulatory grammar compatibility across backgrounds is non-negotiable regardless of how much slack the protein itself has. The framework predicted a firmware layer. The data returned a firmware layer with an internal structure the framework also predicts.

Three independent lines of evidence --- predictive performance, variance compression, GO enrichment --- converge on the same conclusion. That is not proof of causality. The analysis is correlational, the annotations are incomplete, and the mechanistic score captures structure without establishing mechanism. But convergence across three independent tests, each designed to be falsifiable by a different class of confound, is not consistent with artifact. It is consistent with a compatibility layer.

This framework sits upstream of established hypotheses for sex, not in competition with them. Classic models emphasize specific benefits: parasite-mediated selection (Red Queen), purging of deleterious mutations (Muller’s Ratchet). Flexible determinism treats the act of mixing itself as a constraint. Rather than paradox, sex is a filter --- one that continuously converts background-dependence into a selectable liability, preserving the shared molecular language that complex multicellular life requires.

The most rigorous test of this hypothesis requires comparing sexual lineages against obligate asexual sister clades. That test is not executed here, and the reasons are methodological, not theoretical. Standard sexual--asexual comparisons are confounded by two problems. First, genome-wide mutational decay in asexuals (Muller’s ratchet). Second, differences in effective population size. Obligate asexual eukaryotes compound the problem: they are rare, often cryptically sexual, or of hybrid origin --- bdelloid rotifers, parthenogenetic lizards --- making clean inference difficult.

The right design measures the *internal contrast* within each genome: the ratio of firmware-site constraint to bulk synonymous constraint. In sexual lineages, that ratio should be high ---firmware is maintained by population-wide compatibility pressure. In independently derived asexual lineages, it should collapse. Ratios rather than absolute constraint levels control for the population-size and mutational-load differences that sink direct inter-lineage comparisons.

Cyclical parthenogens with living sexual sister lineages --- *Timema* stick insects, *Daphnia* --- are the appropriate comparison units. This is the designated falsification test for this framework.

### Box 2.

#### Logical extensions: predictions the framework generates beyond the data presented here

The compatibility-feedback framework makes several predictions that extend beyond the mammalian analysis presented in this paper. We do not test these here, but we flag them because they follow directly from the core logic --- and because a framework that coheres with independent phenomena is more likely to be capturing something real than one that explains only the data it was built on.

**Incest avoidance.** If the value of sexual reproduction is that genome-matching between unrelated individuals provides valid signal about what is working --- the more two strangers share a sequence feature, the more likely that feature is genuinely useful rather than merely present --- then matching between related individuals degrades that signal. Shared sequence in relatives may reflect common descent rather than independent convergence on a functional solution. Under this framework, incest avoidance is not primarily a genetic-load story, though load effects are real. It is an information-quality story: outcrossing with unrelated individuals maximizes the validity of the compatibility signal. This predicts that incest avoidance should be strongest in lineages where compatibility filtering is most critical --- broadly expressed, core-machinery genes --- and weakest where local adaptation dominates.

**Speciation.** The compatibility signal is only valid within a shared selective environment. Two populations adapting to different ecological contexts --- different predators, different food sources, different thermal regimes --- will develop different firmware requirements. Matching between individuals from those populations is no longer informative about what works in either environment; it is noise. Under this framework, reproductive isolation is not merely a byproduct of divergence. It is the genome’s way of preserving the validity of its own compatibility signal. Speciation is where the feedback loop closes: once environments diverge enough that cross-population matching no longer generates valid signal, selection favors mechanisms that prevent the mixing. This predicts that reproductive isolation should evolve faster in lineages with strong firmware architecture than in lineages with weak firmware architecture, controlling for ecological divergence.

*These extensions are speculative. They are offered as predictions, not findings, and as an invitation for the framework to be tested against independent lines of evidence*.

Two limitations are worth naming. First, the analysis is correlational and relies on existing annotations; causality requires experimental perturbation and more explicit modeling of context-dependent mutability, particularly methylation chemistry --- CpG status is the dominant ablation block, and that fact demands a richer mutation model. Second, the variance compression result, while striking, could reflect unmeasured covariates correlated with the firmware score; future work should test it against expression-breadth and tissue-specificity controls.

The *Drosophila melanogaster* control confirms that regulatory proximity carries a constraint signal outside mammals. However, confirming the phenomenology in a sexual species is not the same as testing the sexual--asexual boundary. If the model holds, asexual lineages --- which lack the cross-individual compatibility filter --- should show systematically weakened mechanistic constraint at synonymous sites. Synonymous sites are not silent observers. They are active participants in the interoperability scaffold that makes sexual reproduction possible.

## Methods

Data source. We used mammal_4d_constraint_analysis.RData (Figshare DOI: 10.6084/m9.figshare.26318410), loaded into R as a data.table with 2,621,118 rows and 72 columns. We analyzed phyloP (phyloP.am), which was numerically identical to phyloP in this dataset (matching summaries).

Variable mapping. Response: y: = phyloP.am. Predictors: gc: = prop_GC; gc1mb: = GC_1mb_window; expr: = mean_expression; mu: = mu_syn; cpg_site: = as.integer(CpG_site.y==1); boundary20: = as.integer(min_boundary_dist<20); ese: = as.integer(in_ESE); ese520: = as.integer(in_ESE_520); tfbs: = as.integer(TFBS_overlap>0); ccre: = as.integer(cCRE_overlap>0); rbp: = as.integer(RBP_overlap>0).

Mutation-rate proxy. The covariate mu (mu_syn in the source table) was used as an estimated synonymous mutability proxy to absorb regional variation in substitution propensity unrelated to functional constraint. In the analysis code, mu was defined as a numeric copy of mu_syn (Analysis script, line 15; Code availability). Across sites with finite mu_syn (n = 2,438,084; DT: 2,621,118 rows with 183,034 missing mu_syn), mu spans 2.49×10⁻⁸ to 3.67×10⁻⁴ (median 1.03×10⁻⁵; mean 1.59×10⁻⁵). mu_syn is only weakly correlated with phyloP (Pearson r = −0.0447; Spearman ρ = −0.0653), arguing against a trivial circular dependence between the proxy and the response. As a robustness check under the strengthened baseline (tri3+codon3), excluding mu reduced held-out R² only modestly (0.3065→0.3036; ΔR² over baseline 0.0933→0.0905; Extended Data Table 2), indicating that the mechanistic signal is not driven by the mutation-rate proxy.

Train/test split. We held out chromosomes chr1, chr10, chr12 and chr15 for evaluation. The training set included all remaining chromosomes. For each model we used complete-case filtering for its required variables; no imputation was used.

Model specification. Baseline model: y ∼ gc + ns(gc1mb, df=5). Full model: y ∼ gc + ns(gc1mb, df=5) + expr + mu + boundary20 + ese + ese520 + tfbs + ccre + rbp + cpg_site. Models were fit by ordinary least squares; held-out performance was measured via R² = 1 − RSS/TSS on the test set.

Mechanistic increment score. On the held-out set we computed pred_base and pred_full. The mechanistic increment score was score: = pred_full − pred_base. The GC-residualized outcome was resid_gc: = y − pred_base. Association metrics were obtained from resid_gc ∼ score and from decile binning of score.

Bootstrap confidence interval. To quantify uncertainty on ΔR², we performed 1,000 bootstrap iterations via chromosome-level resampling of the four held-out chromosomes (with replacement). For each iteration we computed ΔR² = R²(full) − R²(base) on the resampled held-out set. The 95% CI was taken as the 2.5th and 97.5th percentiles of the bootstrap distribution (n_boot = 1,000; ΔR² = 0.129, 95% CI: [0.127, 0.131]).

Variance compression analysis. We merged the 4D site-level dataset with transcript-level mean mechanistic increment scores (n = 3,582 transcripts with ≥10 held-out sites). Transcripts were ranked by mean score and binned into 10 equal-count deciles. Within each decile we computed the mean and standard deviation of phyloP scores. GC-residualized analysis used resid_gc in place of raw phyloP. Statistical significance was assessed via Wilcoxon rank-sum test on decile SD values vs. decile rank.

LOEUF co-conservation analysis. We merged transcript-level firmware scores with gnomAD v2.1.1 LOEUF scores by Ensembl transcript ID (n = 3,452 matched of 3,582). We regressed mean_loeuf ∼ mean_score and mean_pLI ∼ mean_score by ordinary least squares. Decile summaries were computed as above.

GO enrichment analysis. Gene lists were constructed from the top 10% (n = 345) and bottom 10% (n = 345) of transcripts by mean firmware score, and two subsets of the top 10% split by LOEUF tertile (hi_loeuf: n = 172; lo_loeuf: n = 173). Enrichment was performed using g:Profiler (g:GOSt, accessed 2026) with organism = hsapiens, correction_method = g_SCS, and all GO (BP, MF, CC) plus KEGG and Reactome sources. Background was set to all protein-coding transcripts in the dataset.

Ablations. Predictor blocks were defined as CpG = {cpg_site}; ExprMu = {expr, mu}; Boundary = {boundary20}; ESE = {ese, ese520}; Overlap = {tfbs, ccre, rbp}. We refit the model excluding one block at a time and recomputed held-out R², ΔR² relative to baseline, and resid_gc∼score metrics.

Sequence-context and codon-usage robustness. To evaluate whether motif-based predictors merely proxy for unmodeled sequence composition, we strengthened the baseline with two categorical controls: (i) a local 3-mer context factor (tri3) extracted from the central triplet in codonString and restricted to valid A/C/G/T triplets, and (ii) a codon identity factor (codon3) for the synonymous-site triplet. To efficiently estimate these high-cardinality fixed effects, robustness models were fit on a stratified random subsample of 800,000 training sites (balanced across chromosomes). At this sample size, parameter estimates for tri3/codon3 are statistically stable (standard errors negligible), and evaluation remained on the full held-out chromosome set. Software and reproducibility. Analyses were performed in R 4.5.2 with data.table, splines, ggplot2 and pROC. The reproducibility package includes the analysis script, tables and figures, and sessionInfo.

## Data availability

The *D. melanogaster* dm6 reference genome sequence (dm6.fa.gz) and Ensembl/UCSC gene annotations (dm6.ensGene.gtf.gz) were obtained from the UCSC Genome Browser downloads [16] (goldenPath dm6 bigZips). The dm6 phyloP 27-way conservation track (dm6.phyloP27way.bw) was obtained from the UCSC dm6 phyloP download directory. The mammalian dataset used for the primary analyses is publicly available on Figshare (DOI: 10.6084/m9.figshare.26318410). All analysis scripts, derived intermediate files, and manuscript-ready outputs generated in this study are provided in the accompanying reproducibility package.

## Code availability

All code required to reproduce the analyses, including run scripts, session information, and pre-generated tables/figures, is provided in the accompanying reproducibility package.

## Acknowledgements

The author thanks open-source software communities for the tools used in this study.

## Extended Data

### Tables

**Extended Data Table 1.** Leave-one-block-out ablation under strong baseline (trinucleotide context + codon identity).

| model | R2_base | R2_full | dR2 | beta_resid_score | R2_resid_score | delta_resid_per_1SD |
| --- | --- | --- | --- | --- | --- | --- |
| FULL | 0.2131 | 0.3064 | 0.09332 | 1.018 | 0.118335 | 1.12273 |
|  | 59 | 87 | 85 |  |  |  |
| No_ESE | 0.2131 | 0.3063 | 0.09314 | 1.00699 | 0.118098 | 1.12161 |
|  | 59 | 04 | 57 |  |  |  |
| No_ExprMu | 0.2131 | 0.3036 | 0.09044 | 0.995778 | 0.114665 | 1.10519 |
|  | 59 | 04 | 52 |  |  |  |
| No_Boundary | 0.2131 | 0.3014 | 0.08828 | 1.00996 | 0.111939 | 1.09197 |
|  | 59 | 46 | 74 |  |  |  |
| No_Overlap | 0.2131 | 0.3007 | 0.08761 | 1.00087 | 0.111045 | 1.0876 |
|  | 59 | 72 | 28 |  |  |  |
| No_CpG | 0.2131 | 0.2293 | 0.01618 | 1.13574 | 0.0209686 | 0.472612 |
|  | 59 | 49 | 99 |  |  |  |

**Extended Data Table 2.** Sensitivity to removing μ (mutation-rate proxy) under strong baseline (trinucleotide context + codon identity). (A) Predictive performance on held-out chromosomes (R² base, R² full, ΔR²). (B) Residual-to-score scaling after residualizing on the strong baseline (β, R², and residual shift per 1 SD score).

| Model | (A) Predictive performance | (B) Residual-to-score scaling |
| --- | --- | --- |
| | $R^2(\text{base})$ | $\beta(\text{resid} \sim \text{score})$ |
| | $R^2(\text{full})$ | $R^2(\text{resid} \sim \text{score})$ |

| | $\Delta R^2$ | $\Delta$ residual per 1 SD score |
| --- | --- | --- |
| Full model (with $\mu$) | 0.213159 | 1.018 |
|  | 0.306487 | 0.118335 |
|  | 0.0933285 | 1.12273 |
| Full model (without $\mu$) | 0.213159 | 0.995778 |
|  | 0.303604 | 0.114665 |
|  | 0.0904452 | 1.10519 |

**Extended Data Table 3.**
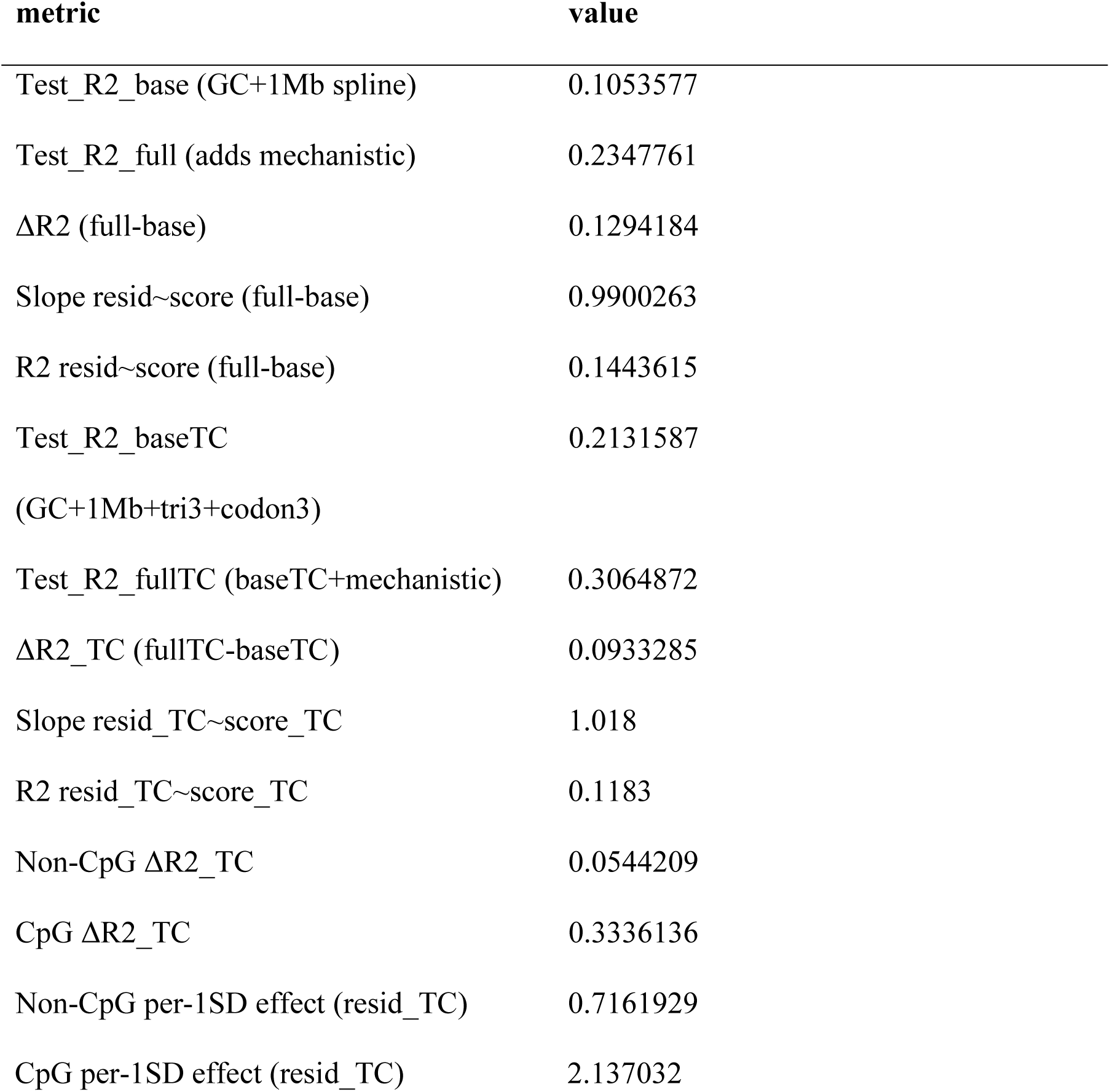
Held-out performance metrics and score–residual association.

**Extended Data Table 4.** Leave-one-block-out ablation results.

| model | R2_base | R2_full | dR2 | beta_resid<br>_score | R2_resid_sc<br>ore | delta_resid_<br>per_1SD |
| --- | --- | --- | --- | --- | --- | --- |
| FULL | 0.10535769 | 0.23477610 | 0.12941840 | 0.99002626 | 0.14436145 | 1.322789501 |
|  | 5682082 | 2059983 | 6377901 | 1458994 | 8333289 | 3461 |
| No_ES | 0.10535769 | 0.23406963 | 0.12871193 | 0.99022917 | 0.14356715 | 1.319145345 |
| E | 5682082 | 3176611 | 7494529 | 2189282 | 049775 | 33313 |
| No_Exp | 0.10535769 | 0.23267486 | 0.12731716 | 0.98349921 | 0.14206126 | 1.312208806 |
| rMu | 5682082 | 4262108 | 8580027 | 6885914 | 5110432 | 31047 |
| No_Bo | 0.10535769 | 0.22838538 | 0.12302768 | 0.99005171 | 0.13721907 | 1.289651453 |
| undary | 5682082 | 4371912 | 868983 | 191517 | 4552653 | 78644 |
| No_Ov | 0.10535769 | 0.22801022 | 0.12265252 | 0.98181904 | 0.13679627 | 1.287663080 |
| erlap | 5682082 | 4092365 | 8410283 | 9148075 | 4383234 | 37138 |
| No_Cp | 0.10535769 | 0.12499004 | 0.01963234 | 1.11542976 | 0.02224537 | 0.519260147 |
| G | 5682082 | 2773593 | 70915113 | 322261 | 68332929 | 730605 |

**Extended Data Table 5.** Headline performance metrics.

| Metric | Value |
| --- | --- |
| Test R<sup>2</sup> base (GC + 1Mb spline) | 0.1054 |
| Test R<sup>2</sup> full (adds mechanistic) | 0.2348 |
| ΔR<sup>2</sup> (full – base) | 0.1294 |
| Bootstrap 95% CI lower | 0.1274 |
| Bootstrap 95% CI upper | 0.1314 |
| Bootstrap n iterations | 1,000 |
| Slope resid~score (full–base) | 0.9900 |
| R<sup>2</sup> resid~score | 0.1444 |
| Test R<sup>2</sup> base TC (GC + 1Mb + tri3 + codon3) | 0.2132 |
| Test R<sup>2</sup> full TC (baseTC + mechanistic) | 0.3065 |
| ΔR<sup>2</sup> TC | 0.0933 |
| Slope resid_TC~score_TC | 1.018 |
| R<sup>2</sup> resid_TC~score_TC | 0.1183 |
| Non-CpG ΔR<sup>2</sup> TC | 0.0544 |
| CpG ΔR<sup>2</sup> TC | 0.3336 |
| Non-CpG per-1SD effect (resid_TC) | 0.716 |
| CpG per-1SD effect (resid_TC) | 2.137 |

**Extended Data Table 6.** Variance compression summary.

| Decile | N transcripts | Mean phyloP | SD phyloP | Mean score |
| --- | --- | --- | --- | --- |
| 1 | 358 | −0.34 | 4.08 | −0.766 |
| 2 | 358 | −0.18 | 3.91 | −0.358 |
| 3 | 358 | 0.02 | 3.75 | −0.179 |
| 4 | 358 | 0.14 | 3.61 | −0.063 |
| 5 | 358 | 0.28 | 3.49 | +0.051 |
| 6 | 358 | 0.41 | 3.35 | +0.151 |
| 7 | 358 | 0.55 | 3.19 | +0.241 |
| 8 | 358 | 0.67 | 3.04 | +0.318 |
| 9 | 358 | 0.79 | 2.86 | +0.399 |
| 10 | 358 | 0.91 | 2.68 | +0.542 |

**Extended Data Table 7.** GO enrichment top terms by gene list.

| List | Top term | GO ID | Adjusted p |
| --- | --- | --- | --- |
| GO_A (high firmware) | Cytosol | GO:0005829 | $6.9 \times 10^{-213}$ |
| GO_A (high firmware) | Cytoplasm | GO:0005737 | $3.5 \times 10^{-212}$ |
| GO_A (high firmware) | Protein binding | GO:0005515 | $7.6 \times 10^{-13}$ |
| GO_B (low firmware) | Cell differentiation | GO:0030154 | $6.5 \times 10^{-11}$ |
| GO_B (low firmware) | Regulation of membrane potential | GO:0042391 | $5.6 \times 10^{-10}$ |
| GO_B (low firmware) | Nervous system development | GO:0007399 | $2.1 \times 10^{-9}$ |
| GO_B (low firmware) | Neurogenesis | GO:0022008 | $6.6 \times 10^{-9}$ |
| GO_C (decoupled) | Mitochondrion | GO:0005739 | $3.1 \times 10^{-6}$ |
| GO_C (decoupled) | Carboxylic acid<br>metabolic process | GO:0019752 | $2.6 \times 10^{-2}$ |

**Extended Data Table 8.** Robustness to strengthened sequence-context baselines.

| model | R2 | dR2_vs_base |
| --- | --- | --- |
| base (gc + ns(gc1mb)) | 0.1044022 | 0.0000000 |
| base+tri3+codon3 | 0.2131587 | 0.1087565 |
| full (adds mechanistic) | 0.2351259 | 0.1307236 |
| full+tri3+codon3 | 0.3064872 | 0.2020850 |

**Extended Data Table 9.**
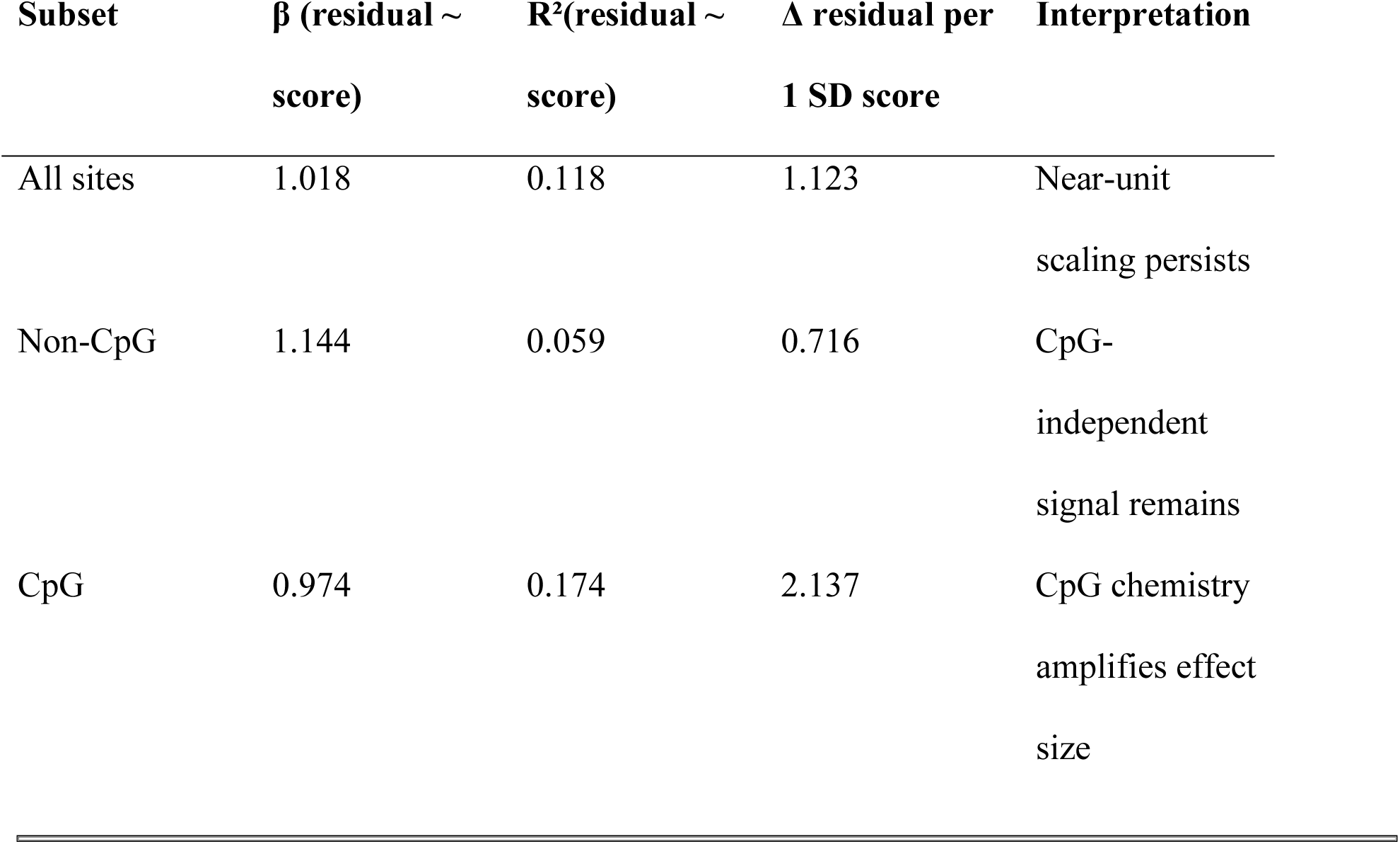
Residual-to-score scaling after residualizing on the strengthened baseline (trinucleotide context ± codon identity).

## Supplementary Discussion

Positioning relative to existing theories of sex. As outlined in the Discussion, the compatibility-feedback framing overlaps with several well-studied ideas but differs in emphasis. Traditional population-genetic accounts treat sex primarily as a mechanism for generating and combining variation (recombination) and for managing linkage disequilibrium under selection. The compatibility lens instead treats the act of mixing as an informational constraint: outcrossing repeatedly exposes components to heterogeneous partner contexts, turning background dependence into a selectable liability. In that sense, compatibility feedback is a ‘selection on robustness’ interpretation rather than a ‘selection on variation generation’ interpretation. These are compatible: recombination both generates novel combinations and reveals which components remain viable across them.

This section expands conceptual interpretation and robustness considerations, and is intended as online-only discussion rather than the core empirical narrative.

Relationship to major hypotheses for sex and recombination. The compatibility-feedback framework sits upstream of several established hypotheses for the maintenance of sex, treating them as context-specific consequences of a broader requirement: maintaining a stable functional substrate while permitting open-ended phenotypic exploration ([5–7, 17–23]).

Parasite-mediated selection and Red Queen dynamics. Classic parasite/host coevolution models explain sex as a strategy for continually generating novel genotypes in response to rapidly evolving antagonists. Under flexible determinism, such dynamics are expected in any system where intimate contact and horizontal microbial exposure are recurrent features of reproduction, so Red-Queen pressure is treated as a powerful selective context rather than the root driver of sex ([3, 4]).

Mutation-purging, Muller’s ratchet, and synergistic epistasis. Models emphasizing the removal of deleterious mutations (including ratchet-type arguments) are naturally compatible with an n=2 feedback interpretation: comparison and recombination between unrelated genomes provides a mechanism to prevent irreversible mutational accumulation and to preferentially retain shared, high-utility sequence states. In this view, “mutation purging” is not an alternative explanation but a concrete, falsifiable consequence of stronger stabilizing selection at the most interoperable loci ([5, 6, 20]).

Recombination-accelerated adaptation (Fisher–Muller class). Another major family of arguments treats recombination as a way to combine beneficial variants and reduce clonal interference, accelerating adaptation. Flexible determinism is consistent with this while emphasizing a complementary role: recombination can both assemble advantageous combinations and sharpen which components become “firmware-like” through recurrent cross-individual compatibility ([6, 21, 22]).

DNA repair / meiotic repair hypotheses. The observation that meiosis includes repair-like machinery has motivated proposals that the primary role of sex is DNA repair. Here, repair is treated as a mechanistic implementation detail of recombination rather than a sufficient rationale by itself: repair maintains structure, but the framing asks what selects for maintaining particular structures. Compatibility-feedback provides an explicit selection-level motive for maintaining, and preferentially stabilizing, shared functional sequence states ([23]).

Ecological and diversity-maintenance hypotheses (tangled bank; bet-hedging). Ecological models propose that sex is favored because offspring diversity reduces sibling competition or spreads risk across heterogeneous environments. These are downstream consequences: once a stable substrate is maintained, sex generates diverse phenotypic applications adapted to local niches. ([7]).

Sexual selection and mate choice. Sexual selection can modulate which variants enter the recombining pool, but it presupposes a sex-enabled mating system; under the present framing, mate choice is a higher-level filter operating on top of the stability/variation tradeoff created by sex, not the origin of that tradeoff.

Meiotic repair as mechanism vs rationale. “DNA repair” in meiosis is often described as a reason for sex, but repair is intrinsically relational: it maintains structure in service of an objective. In this framing, the objective is the stabilization of shared functional “firmware” across unrelated genomes, with repair machinery providing one way to implement recombination and maintain genome integrity during that process ([22]).

Mutation purging as a falsifiable consequence. If stabilizing selection is stronger at loci that repeatedly pass the cross-individual compatibility filter, then mutation density (and tolerated functional variation) should be lower in those loci, while higher-variance loci should show more standing variation. This yields a qualitative, testable prediction: within sexually reproducing lineages, regions with stronger compatibility-feedback signatures should show reduced tolerated divergence and reduced accumulative mutational burden, controlling for local mutation processes ([17, 19]).

CpG stratification and interpretation. CpG context is the dominant single block in the ablation table, reflecting both mutation-rate heterogeneity and selection. However, within CpG and non-CpG strata, the mechanistic score remains associated with phyloP residuals, suggesting that the model is not merely learning CpG. One practical implication is that future modeling could explicitly parameterize mutation processes (e.g., context-dependent mutation rates) and then test whether mechanistic annotations retain incremental value under a richer mutation model.

Speciation and inbreeding avoidance as compatibility-information hypotheses. The notion that inbreeding reduces the informativeness of the compatibility signal can be expressed in journal terms as an information-gain argument about genotype–genotype interaction screening.

Likewise, the idea that reproductive isolation can preserve the meaning of compatibility signals across ecological regimes can be stated as a hypothesis about selection maintaining coherence between constraint landscapes and recombination pools.

Note on synonymous-site selection. Synonymous positions can be targets of selection via regulatory/splicing effects and other mechanisms; this matters because phylogenetic conservation can reflect multiple selective regimes rather than a single cause [24], and synonymous patterns can also reflect codon-usage and translational selection [25, 26] or adaptive dynamics with different sweep modes [27].

## Supplementary Results: Cross-taxon counterfactual in Drosophila melanogaster

Rationale. A core vulnerability of any argument linking synonymous-site constraint to sexual reproduction is the absence of an explicit counterfactual. As an initial cross-taxon test, we evaluated whether the same type of mechanistic signal seen in mammals generalizes to Drosophila melanogaster (dm6). This does not test the sexual–asexual boundary directly; rather, it tests whether the mammalian signal is a parochial artifact of mammalian sequence context, annotation, or CpG chemistry.

Data and site definition. We downloaded the dm6 reference genome (FASTA), gene annotation (GTF), and a multi-species phyloP conservation track (27-way) and constructed a catalogue of fourfold-degenerate (4D) synonymous sites from annotated coding sequences. For each unique genomic site we retained (i) local 3-mer context, (ii) codon identity at the site (codon3), (iii) the number of transcripts sharing the site (n_tx), and (iv) distance to the nearest exon boundary (min_boundary_dist). Conservation was quantified as the phyloP score at that position.

Models. To parallel Track 1, we used a conservative baseline that controls for sequence context and codon identity: y ∼ trinucleotide context + codon identity + n_tx (base0). We then added a flexible boundary-distance term using a spline on log-transformed distance: y ∼ trinucleotide context + codon identity + n_tx + ns(log_dist, df=5) (+dist). We also evaluated a simple boundary indicator (boundary20) alone and in combination with the distance spline. Models were trained on a chromosome-holdout split and evaluated on held-out chromosomes.

Results. In dm6, adding the boundary-distance spline improved held-out prediction relative to the context-only baseline (base0 R² = 0.0417; +dist R² = 0.0705; ΔR² = +0.0288). In contrast, a simple boundary20 threshold did not add predictive power beyond the baseline (base0 R² = 0.0417; +boundary20 R² = 0.0565; +dist+boundary20 R² = 0.0706; ΔR² vs +dist = +7.8×10⁻⁵). The increment score from the +dist model (pred_dist − pred_base0) predicted residual conservation with near-unit scaling (slope = 0.995; R²(resid∼score) = 0.0300), and decile binning showed a monotone trend from from strongly negative to strongly positive residuals (Supplementary Fig. 1); mean −0.16 to +0.43, standard error ∼0.002.”. These results support a conserved *phenomenology*—sequence-context–controlled regulatory proximity carries information about evolutionary constraint—while also indicating that the most informative boundary representation differs between mammals (boundary threshold is salient) and flies (continuous distance is salient). A plausible interpretation is architectural: mammalian exon-definition may impose relatively sharp, motif-dense boundary zones, whereas Drosophila intron-definition can yield a broader, distance-dependent gradient of constraint.

Hexamer context control (additional baseline hardening). To stress-test the concern that distance-to-boundary effects may proxy for unmodeled local k-mer composition, we added a 6-mer context control (hex6) computed from the reference sequence at each site. Because hex6 has high cardinality, the hex6-augmented models were fit on a large stratified subsample of the training data, then evaluated on the full held-out set. Adding hex6 increased held-out performance beyond the +dist model (R²(+dist) = 0.0678; R²(+dist+hex6) = 0.0853; ΔR² = +0.0175).

Residualization against the +dist model showed that the hex6 increment (pred_dist+hex6 − pred_dist) remained systematically associated with residual conservation (slope = 0.690; R² = 0.0237; n_used = 357,383; n_dropped_missing = 195). Together, these controls indicate that local sequence context can explain additional variance, but does not eliminate the boundary-distance signal; rather, both contribute.

We included a 6-mer (hex6) control only in Drosophila because the fly analysis is orders of magnitude smaller than the mammalian dataset, making higher-order k-mer fixed effects computationally tractable. In mammals, the strengthened trinucleotide context + codon identity baseline already absorbs the dominant codon-level and immediate-context effects and is intentionally conservative; adding hexamer controls at mammalian scale would be substantially more expensive and would likely further reduce the apparent incremental contribution of annotation-derived predictors. Accordingly, mammalian ΔR² estimates under the strengthened baseline should be interpreted as lower bounds on the predictable, mechanism-linked component beyond simple sequence context.

Interpretation for the sexual-feedback hypothesis. In dm6, adding the boundary-distance spline improved held-out prediction relative to the context-only baseline (base0 R² = 0.0417; +dist R² = 0.0705; ΔR² = +0.0288). The increment score from the +dist model predicted residual conservation with near-unit scaling (slope = 0.995; R²(resid∼score) = 0.0300), and decile binning showed a monotone trend from strongly negative to strongly positive residuals. These results support a conserved phenomenology — regulatory proximity carries information about evolutionary constraint across phylogenetically distant animal genomes — while indicating that the most informative boundary representation differs between mammals (threshold is salient) and flies (continuous distance is salient). Mammalian exon-definition imposes sharp, motif-dense boundary zones, whereas Drosophila intron-definition yields a broader, distance-dependent gradient.

Limitations. The present analysis uses phyloP as a proxy for long-term constraint and focuses on 4D sites to avoid amino-acid changes, but phyloP integrates multiple evolutionary processes and is not a direct readout of “compatibility.” Moreover, annotations such as TFBS/cCRE/RBP overlaps are incomplete and context-dependent. The mechanistic model should therefore be read as an empirical scaffold: evidence that interpretable features explain a robust component of conservation beyond composition, rather than a complete or final model of constraint.

**Supplementary Fig. 1.**
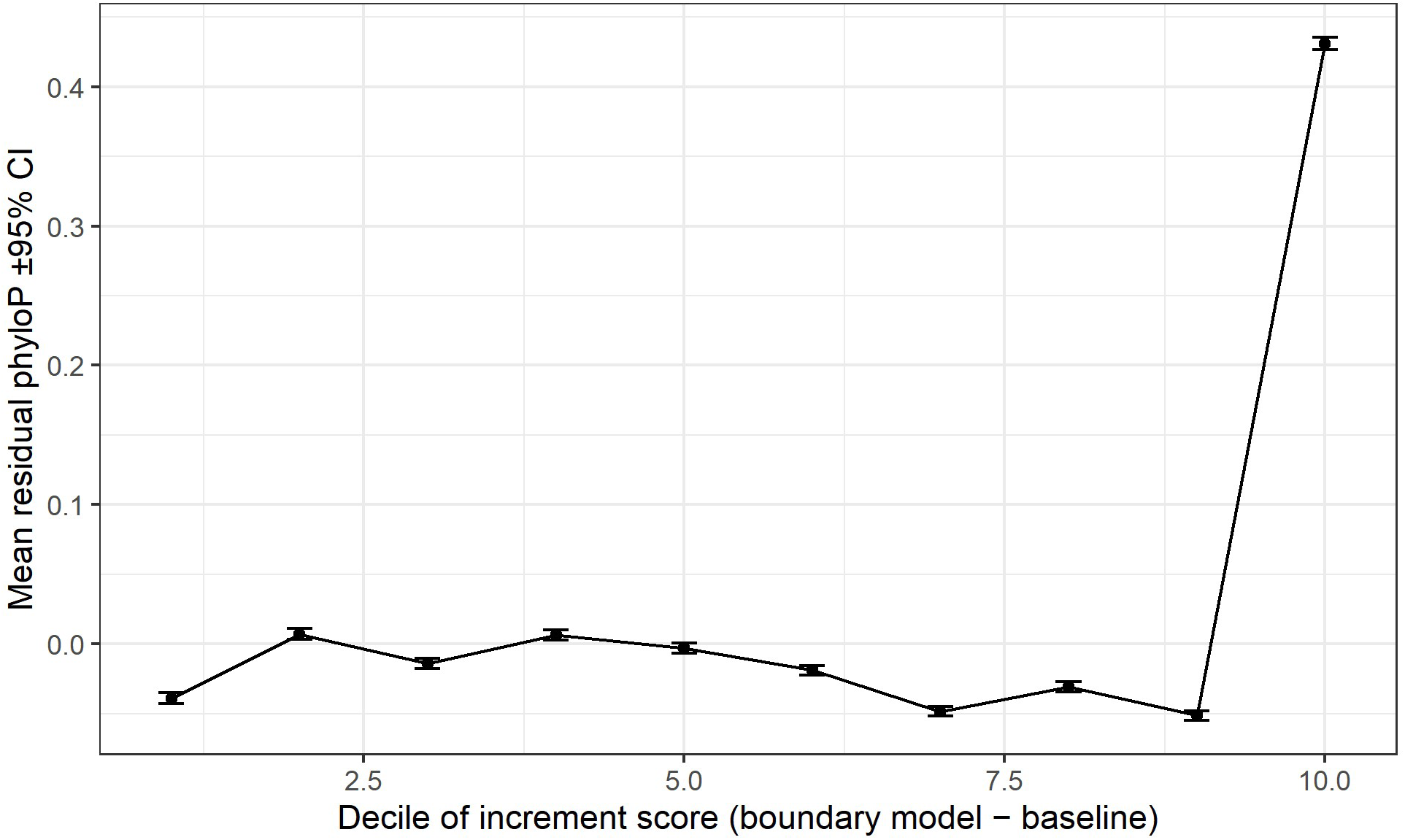
Validation of mechanistic constraint in *Drosophila melanogaster*. Synonymous sites in *D. melanogaster* (dm6) were binned by proximity to splice boundaries (deciles), and the mean residual conservation score (PhyloP) was calculated for each bin after controlling for local trinucleotide and codon context. The monotonic positive trend confirms that regulatory proximity predicts evolutionary constraint beyond sequence background in fruit flies, paralleling the mammalian result. Source data are provided in Track2_Table_BoundaryDist_Deciles.tsv.

## Supplementary Methods

Alternative baselines and why GC+gc1mb was chosen. GC is correlated with substitution processes and with many genomic features. A baseline using only site-level GC can be too weak, inflating apparent contributions of other features that correlate with regional GC. Conversely, overly complex baselines can absorb biologically meaningful signal. We used site-level GC plus a flexible spline in 1 Mb GC as a conservative middle ground. The spline captures non-linear regional effects while remaining interpretable.

Potential extensions: generalized additive models and interactions. The present linear framework intentionally prioritizes interpretability. However, functional constraints may involve interactions: for example, splice proximity might matter more in highly expressed genes, or CpG effects might differ near boundaries. Generalized additive models or tree-based models can capture such interactions. A reviewer-proof escalation is to report that linear models already capture substantial signal on held-out chromosomes, and to position non-linear models as future work (or as a Supplementary analysis).

Block definition rationale for ablations. Blocks were chosen to group conceptually related predictors that might jointly represent a mechanistic channel: (i) CpG as methylation/mutation context; (ii) ExprMu as expression-linked constraint and mutation propensity; (iii) Boundary as splice-adjacent constraint; (iv) ESE as explicit splicing-enhancer overlap; (v) Overlap as regulatory/RNA-binding overlaps. Ablations are not causal identification; they approximate ‘importance’ under collinearity.

Decile plots and uncertainty. Decile binning of score provides a non-parametric view of monotonic trends. Because the dataset is large, standard errors on binned means are small. Confidence intervals shown as ±1.96×SE reflect uncertainty in the mean residual within each bin, not uncertainty in the underlying model parameters.

Why chromosome-level holdout. Whole-chromosome holdout reduces leakage from local correlation and ensures that models generalize across large-scale genomic contexts. A stricter alternative is block holdout using non-overlapping windows genome-wide; this can be added as a robustness analysis if reviewers are concerned that chromosome holdout may still share broad-scale gene-class composition across chromosomes.

